# Surface adhesion and membrane fluctuations influence the elastic modulus of extracellular vesicles

**DOI:** 10.1101/2024.02.07.578591

**Authors:** Fredrik Stridfeldt, Hanna Kylhammar, Prattakorn Metem, Vikash Pandey, Vipin Agrawal, Andre Görgens, Doste R. Mamand, Oskar Gustafsson, Samir El Andaloussi, Dhrubaditya Mitra, Apurba Dev

**Affiliations:** Department of Applied Physics, KTH Royal Institute of Technology, Stockholm, Sweden; Division of Applied Electrochemistry, KTH Royal Institute of Technology, Stockholm, Sweden; Nordita, KTH Royal Institute of Technology and Stockholm University, Stockholm, Sweden; Department of Physics, Stockholm University, Stockholm, Sweden; Department of Materials Science and Engineering, Northwestern University, Evanston, IL 60208; Department of Laboratory Medicine, Division of Biomolecular and Cellular Medicine, Karolinska Institutet, Stockholm, Sweden; Department of Cellular Therapy and Allogeneic Stem Cell Transplantation (CAST), Karolinska University Hospital Huddinge and Karolinska Comprehensive Cancer Center, Stockholm, Sweden; Institute for Transfusion Medicine, University Hospital Essen, University of Duisburg-Essen, Essen, Germany; Department of Electrical Engineering, Uppsala University, Uppsala, Sweden

## Abstract

Elastic properties of nanoscale extracellular vesicles (EVs) are believed to influence their cellular interactions, thus having a profound implication in intercellular communication. Yet, an accurate quantification of the elasticity of such small lipid vesicles is difficult even with AFM-based nanoindentation experiments as it crucially depends on the reliability of the theoretical interpretation of such measurements. Here we describe a complete method composed of theoretical framework, experimental procedure, and appropriate statistical approach for an accurate determination of bending modulus and effective elastic modulus of EVs. Further, we experimentally demonstrate that the quantification of EVs by the elastic modulus from AFM-based force spectroscopy measurement is marred by the interplay of their compositionally inhomogeneous fluid membrane with the adhesion forces from the substrate and thermal effects - two exquisite phenomena that could thus far only be theoretically predicted. The effects result in a large spreading of elastic modulus even for a single EV. Our unified model is then applied to genetically engineered classes of EVs to understand how the alterations in tetraspanin expression may influence their elastic modulus.

## I. INTRODUCTION

The discovery of EVs and their numerous roles in patho-physiological processes are in many ways redefining our approaches and understanding of human health and diseases[1, 2]. Released to the extracellular milieu by most cell types, these nanovesicles carry a wide variety of cellular components including proteins, nucleic acids, and lipids, and are known to deliver their bioactive cargo to a recipient cell, thereby, inducing functional alterations [3]. During the past decade, this topic has generated exploding interest, consequently driving countless technical innovations. While these efforts have helped to overcome many of the initial challenges and skepticism, the high heterogeneity of EVs even when they are released by a single cell has triggered the obvious questions; are there EV subpopulations in circulation that are functionally differential or more relevant than others? If so, which property is the best indicator of such functional differences?

While a single EV approach is arguably the best way to address such heterogeneity, given their nanoscale dimension and diverse molecular repertoire, existing techniques can not perform a comprehensive chemical analysis at a single vesicle. Thus far, optical/fluorescence techniques have been the main methods to interrogate single EVs, although providing only a limited set of information [4, 5]. In this context, the recent attempts made with atomic force microscopy (AFM) has brought renewed enthusiasm as it can provide complementary information, e.g., stiffness, bending modulus, and adhesion [6]. Given the stiffness of EVs may influence their cellular uptake, the topic has also larger implications concerning EV biology and functions [7].

However, being enclosed with a compositionally inhomogeneous fluid-like lipid bilayer, EVs offer yet unexplored challenges for nanoindentation-based measurements, as well as avenues to study some fundamental nature of lipid bilayer which could only be predicted from theory so far. Two such effects are also likely to have significant influence on the membrane elasticity, namely: (i) the competition between adhesion and shape deformation leading to a different mechanism of tether formation for surface adhered vesicles [8, 9], which is the case for most EV studies with AFM and (ii) the effect of membrane fluctuations on the elastic property [10]. To what extent do these effects influence the elastic modulus of EVs remain unknown and so is the accuracy of the previous attempts [11, 12] to quantify the elastic modulus of EVs. Besides, it is also unknown if the alteration in the membrane protein composition can induce measurable changes in EV elasticity.

Here we apply the Canham-Helfrich model (CH model), which has been extensively used to understand force spectroscopy of Giant Unilamellar Vesicles (GUVs) [13], to quantify the elastic properties of EVs from force spectroscopy. We find that in contrast to previous reports [11], the EVs behave as surface-adhered vesicles showing a linear dependence of tether force on the tether length. We then apply our method to different EV populations derived from the human embryonic kidney (HEK293T) cell line and its genetically engineered versions with altered tetraspanin expression. The bending modulus estimated for the different EV families range from 15.7 − 33.0 k_B_T with large fluctuations within each family of vesicles which is consistent with inherent EV heterogeneity. Interestingly, our experiments also show large fluctuations of measured values even for a *single EV* when repeatedly examined under identical conditions. We suggest that such behavior is driven by both thermal and compositional effects, as expected for such nanoscale objects. We find that the probability distribution functions (PDFs) of elastic moduli are best described by not a single Gaussian but superposition of Gaussians. Consequently, commonly used techniques to test for statistical significance, e.g., the Student’s t-test can not be employed. We use Jensen-Shannon divergence to differentiate between the PDFs. Our results suggest that protein alternations might influence the elastic moduli of EVs, but they are often hidden in such large fluctuations.

## II. RESULTS

We performed force spectroscopy in a liquid environment (1× PBS) on EVs derived from HEK293T cell line. We followed the isolation method as described in our previous article [14]. To assess the influence of the protein alterations on the elastic properties of EVs, the expression of tetraspanins was genetically altered in the cell producing two different classes of EVs along with the wild type (WT-EVs): (i) sample with CD63 knockout (CD63-KO) and (ii) sample with CD9-CD63-CD81 knock-out (Pan-KO). To assess the effect of protein corona, if any, we analyzed an additional sample type referred to as WT-SEC. To remove the protein corona, we followed an earlier report [15], and further processed WT-EVs with bind-elution size exclusion chromatography. For more details regarding the sample preparation, see IV C.

### A. Shape of adhered vesicles

A typical topographical profile for an EV in a liquid environment measured by AFM is shown in Fig. (1A). Theories and experiments of adhesion of GUVs to glass cover slips [16–18] suggest that the shape of the vesicle would be that of a spherical cap. Earlier works [6, 19] on nano-vesicles have also made the same assumption. The topographical profile is a height field *H* as a function of two-dimensional coordinates *x* and *y*, typical examples are shown in Fig. (1B). The height field as a function of the slow axis of the scanning of the AFM – along the lines marked in Fig. (1B) – is shown in Fig. (1C). We fit a circle to this profile taking into account the correction due to the shape of the tip of the AFM to determine the effective radius (*R*_c_) of the EV, as earlier described in Ref. [19]. We show in Fig. (1C) the measured height, its fitted arc, and the tip-corrected EV shape for a typical example. The probability distribution of the ratio *H/R*_c_ for all the vesicles is presented in Fig. (1D). A vesicle with *H/R*_c_ *>* 2 indicates a non-realistic shape and was disregarded from future analysis. The radius *R*_c_ ranges from approximately 40 nm to 100 nm. Almost all of them fall within the class of small EVs (sEVs) defined [see e.g., 20, page 9] to be less than 200 nm in diameter but few are not. Hence in the rest of this paper, we call them EVs.

**FIG. 1.**
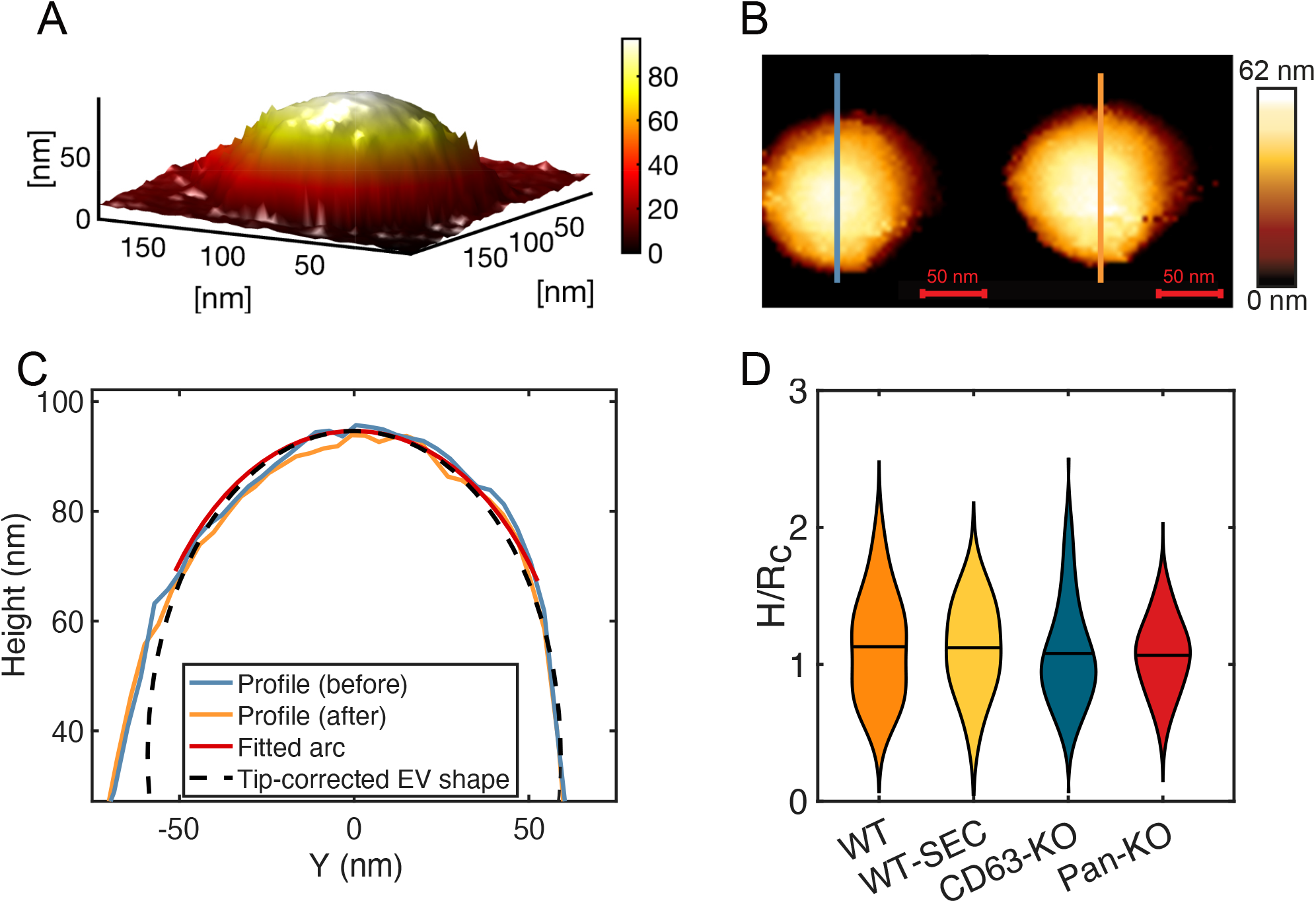
Topographic image. (A) Topographic image of a single vesicle. (B) The height field plotted as a color for a typical vesicle before (left) and after (right) force spectroscopy. (C) The measured height, its fitted arc, and the tip-corrected EV shape extracted from the line shown in (B). (D) The ratio of the height and the radius of curvature. The horizontal black line indicates mean of the ratio. Mean for WT: 1.13, WT-SEC:: 1.12, CD63-KO: 1.08, and Pan-KO: 1.07.

For the measurement of force–distance curves, care is necessary. The EVs need to be immobile for the entire process, and the applied force and tip velocity must be small enough to avoid irreversible damage to the EVs. We image every EV twice – before and after the force spectroscopy – to make sure they are not damaged or have moved, see Fig. (1B) and Fig. (1C). In total, we imaged and performed force-spectroscopy on 49 vesicles of the WT family 39 of the WT-SEC family, 24 of CD63-KO family, and 18 of the Pan-KO family.

### B. Force–distance measurements

In Fig. (2A) we show a typical example of the force– distance curves. Here we plot only the approach curves. As we repeat the experiment many times (≳ 150) on the same EV, each time we obtain a different force–distance curve. This large variation in the force–distance curves is not seen in GUVs. Note that this variation is not originating from the AFM setup, as can be seen by repeating the same experiment but on a clean coverslip instead of an EV, see (Appendix A). We shall revisit these variations later. The average of all the force–distance curves is shown in red in Fig. (2A). We shall call this the *average* force–distance curve.

**FIG. 2.**
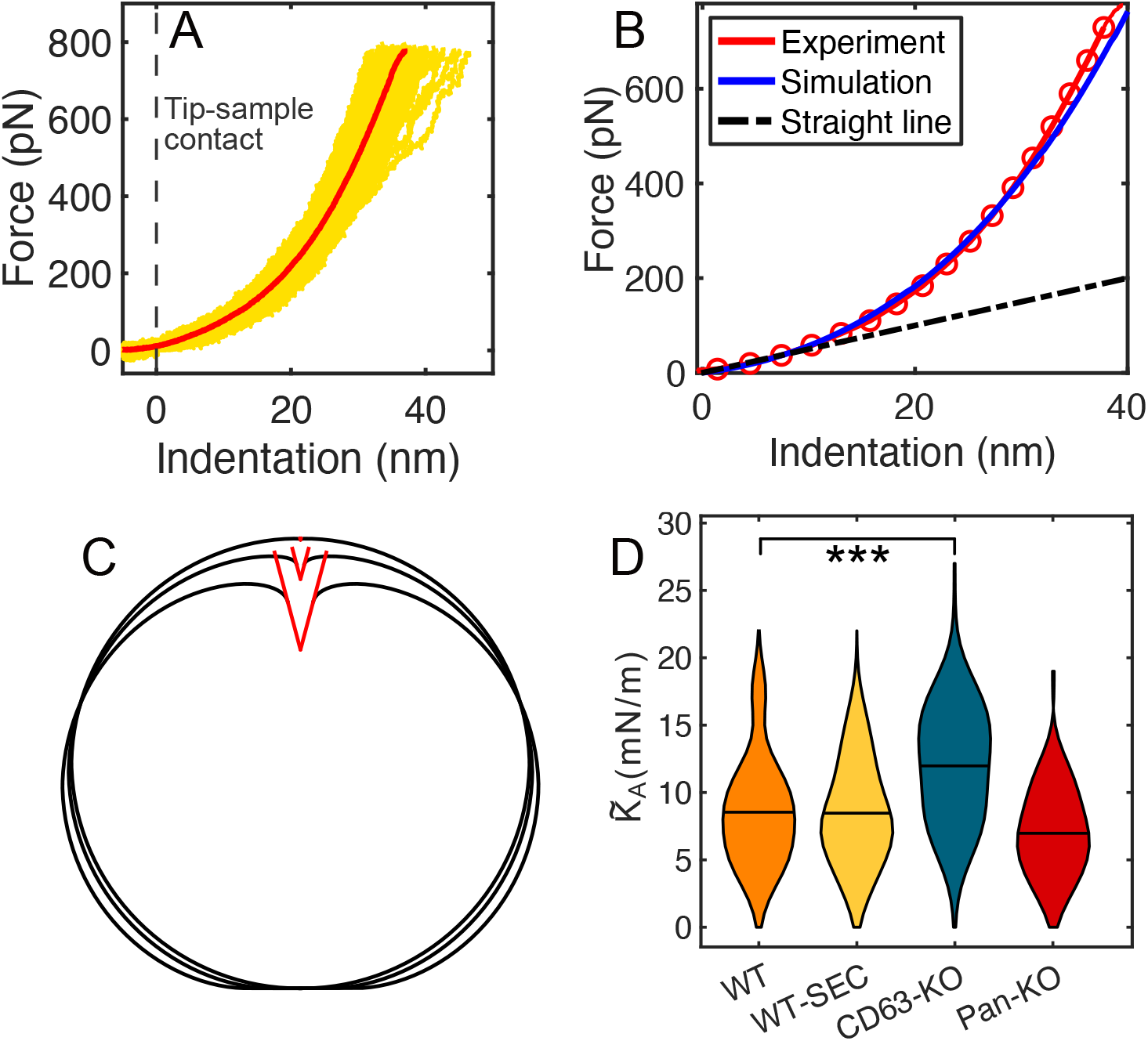
Approach curves. (A) 150 indentation iterations on a vesicle (yellow lines). The average force–distance curve is shown in red. (B) The average indentation curve from (A) and the curve subject to the fitting of the model for *K*_A_ = 618.84 mN m^−1^ and Σ_0_ = 5.49 mN m^−1^ in the same scale. The dashed black line shows a linear fit at small distances. (C) The shape of the vesicle obtained from our simulations for three different values of the force *F* = 2, 90, and 600 pN respectively. The red triangle illustrates the conical tip with the half-cone angle of 15°. (D) The probability distribution functions, plotted as violin plots, of the 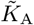 of vesicles from the different families: WT, WT-SEC, CD63-KO, and Pan-KO. These are calculated from the *average* 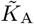 value for each vesicle. The horizontal black line indicates the mean.

#### 1. Theory

The EVs are similar to Giant Unilamellar Vesicles (GUVs) in the sense that both are a drop of fluid enclosed within a lipid bilayer[21]. The two essential differences are: (i) the membrane of the EVs contain many other molecules, including proteins and cholesterol; and (ii) the EVs are much smaller in size. Hence we expect that the theory that describes the force–distance curves of EVs is similar to the theory applied to GUVs. Next, we present a summary of our theory – a more detailed description can be found in (Appendix B).

1. Typically, the force–distance experiments on cells, e.g., red blood cells (RBCs) are interpreted with Hertzian contact mechanics [22–25]. Within this model, the cell is a deformable *solid*. At small distance *d*, the force *F* is given by *F* ∝ *d*^3*/*2^. The EVs cannot be modeled as solids because they lack the internal organelles and cytoskeleton of the cell.
2. The lipid bilayer (with the addition of proteins, nucleic acids, and glycans) of EVs is a (two-dimensional) fluid. It has zero shear modulus but a non-zero area modulus *K*_A_. The usual thin–shell theory [22, 26, 27], which has also been generalized to pressurized shells [10, 28], includes (in–plane) shear modulus and is therefore not applicable to EVs. In other words, the shell in standard thin– shell–theory is a two–dimensional solid whereas the bilipid membrane of EVs is a two–dimensional fluid.
3. Thus, the EVs must be modeled as shells but with fluid membranes, similar to the model of GUVs. We call this the Canham-Helfrich model. In this model, a deformation contributes to the change in energy in three possible ways: change in area, change in volume, and change in curvature or bending. Henceforth we use this model to interpret the force–distance curves.

Assuming that a reasonable estimate of the bending modulus of EVs is the typical bending modulus of lipid bilayers, it is possible to show that the contribution of bending to the change in curvature is very small [6]. Following the standard prescription for GUVs [13] we assume that the volume of the vesicle remains unchanged. This is justified because the only way the volume can change is if water leaks out of the semi-permeable membrane. Using existing data for the permeability of water through bilipid membranes, we find that within the time scale of the experiment the leakage is negligible – the volume remains constant, see (Appendix B 3). Then the only possibility that remains is the change in area.

Let us consider the case where a small external force *F* generates a small deformation *d*. Let *R*_v_ be the effective radius of the vesicle defined by 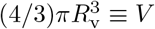, where *V* is the volume of the adhered vesicle. We consider *d* ≪ *R*_v_ such that *d/R*_v_ is a small parameter. Then the change in free–energy can be estimated as (Appendix B 1)

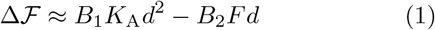

where *B*_1_ and *B*_2_ are dimensionless constants that depend on the details of the indenter and the shape of the adhered vesicle and *K*_A_ is the in–plane area modulus of the membrane. The second term in the free energy is the work done by the indenting force. We have also assumed that the *d* is so small that it does not significantly change the adhesion energy of the EV to the bottom plate. Minimizing the free–energy with respect to *d* and setting it to zero we obtain *F* ∝ *K*_A_*d* – a linear force–distance relationship. Note that thin–shell–theory for pressurized shells also gives a linear force–distance relationship for small deformation [10, 28].

Unfortunately, the linear force–distance relationship for small deformation does not allow us to determine the elastic modulus *K*_A_ because the constant of proportionality, which depends on the adhered shape of the vesicle and the geometry of the indenter, can not be determined by the kind of dimensional arguments given here. Nevertheless, two vesicles with exactly the same surface molecules would have exactly the same interaction with the coverslip and in all cases we have used the AFM probe with the same geometry. Therefore, we can define an effective elastic modulus 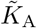 from the slope of the force–distance curve at small deformation. Although this approach does not allow us to measure the *K*_A_ accurately, it does allow us to measure *the change* in *K*_A_ as a result of knocking out of certain proteins. In Fig. (2B) we show a typical example of the average force–distance curve (red line) together with a linear fit at small distances. The slope of the straight line gives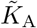.

To fit the force–distance curve over a larger range we need detailed numerical simulations. We follow Ref. [29] who solved the same problem for GUVs, see also (Appendix B 2). Again the only free–energy cost that comes from deformation is due to change in area – the volume is assumed to be constant. The adherence of the vesicle is modeled by a pre–stress, Σ_0_. By obtaining the best fit of the numerical solution of the model to the experimental data we can, in principle, find the two fit parameters, *K*_A_ and Σ_0_. A typical example is shown in Fig. (2B). The numerical and the experimental force–distance curves agree well with each other not only at short distances but over almost the complete range of distances. This confirms that our choice of modeling is appropriate.

#### 2. Experiments

Returning to the experiments we plot the *average* force– distance curve for a single vesicle in Fig. (2B). We find that there is indeed a small range of scale over which *F* ∝ *d*. As the indentation progresses the response becomes nonlinear. Next, we fit a straight line to the first 10% of the indentation (roughly equal to 50 data points) of every indentation’s force–distance curve. The slope of the straight line gives the effective modulus 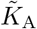 for each vesicle. The probability distribution functions of the average effective modulus 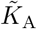 of the families are plotted in Fig. (2D) as violin plots. The mean and standard deviation of the 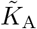 are 8.5 mN*/*m and 3.9 mN*/*m for the WT family, 8.5 mN*/*m and 3.7 mN*/*m for WT-SEC, 12.0 mN*/*m and 3.8 mN*/*m for CD63-KO, and 7.0 mN*/*m and 2.8 mN*/*m for Pan-KO, respectively. In comparison to the WT family, vesicles in the CD63-KO family have a higher mean 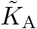 while the Pan-KO family shows a lower mean 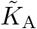. We test the significance of these differences using unpaired two-sample t-tests and find *P*≈ 5 × 10^−4^ when comparing the means of the WT and CD63-KO families. No other family shows a statistical deviation in the mean when compared to the WT family (*P >* 0.05). We shall question the validity of this statistical method in the following section. Earlier studies [30–32] have reported similar effective elastic moduli of various EVs ranging from 7 mN*/*m up to 49 mN*/*m. These studies have all extracted the moduli in a similar way and on similar sample sizes. We have not found any earlier nanomechanical studies on EVs derived from HEK293T cells. To the best of our knowledge the influence of knocking out certain proteins has not been studied before.

The nature of the PDFs plotted in Fig. (2D) suggests that they cannot be approximated by a single Gaussian. This is confirmed by the negative result of the Shapiro– Wilk test, see (Appendix C). We calculate the PDFs using kernel density estimation with a Gaussian kernel. In other words, we find the superposition of Gaussian distributions that best fit our data. In (Appendix D), we plot both the histograms and the PDFs estimated by the kernel density estimation for all the families.

### C. Variation of effective elastic modulus

The PDFs show the inherent heterogeneity of the deformability of the EVs coming from all the different sample families. While the WT, WT-SEC, and Pan-KO families resemble each other in distribution shape, peak, and mean, the CD63-KO distribution shows deviations from the rest. Are the differences we see statistically significant? As the PDFs are not simple Gaussians, the Student’s t-test can not be employed here to test the statistical significance [33]. Hence we need to use a technique that does not assume simple Gaussianity. Furthermore, note that we are dealing with small sample sizes in each family, hence the PDFs themselves are poorly determined. Next, remember that the effective elastic modulus 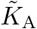 was determined by a linear fit to the *average of all force–distance curves* of a single vesicle but the force–distance curves of a single vesicle show large variation. Thus there are additional sources of error to be taken into account. Let us fit a straight line (at small distances) to the individual force–distance curves for a single vesicle to calculate a 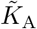for each force–distance curve. Hence for most single vesicles, we obtain about 150 values of 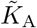. We plot them in Fig. (3A). The values are close to normally distributed but show significant variation, see also (Appendix A). Note that these values are decorrelated with one other. In Fig. (3B) we present the range of values of 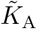that we obtain from individual force–distance curves for each of the families in barcode plots. The families WT and WT-SEC are close to each other although the former shows a larger span of 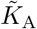 values. The family CD63-KO has relatively high values of 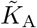 compared to the WT and the family Pan-KO seems to have relatively smaller values of 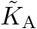 compared to WT. At this stage, we consider for each family three PDFs of 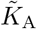 in the following manner. Let the average 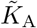 for a single vesicle be denoted by 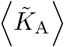, and its standard deviation 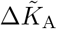. Then we have one PDF each for 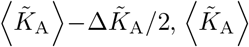, and 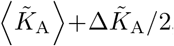, we denote them respectively by 𝒫_1_, 𝒫_2_ and 𝒫_3_. In Fig. (3C) we show that for the WT family. A way to measure the distance between any two PDFs, *p*(*x*) and *q*(*x*) is to calculate their Jensen–Shanon divergence [34] defined by

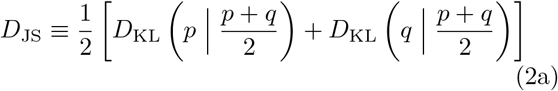

where

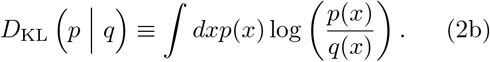

**FIG. 3.**
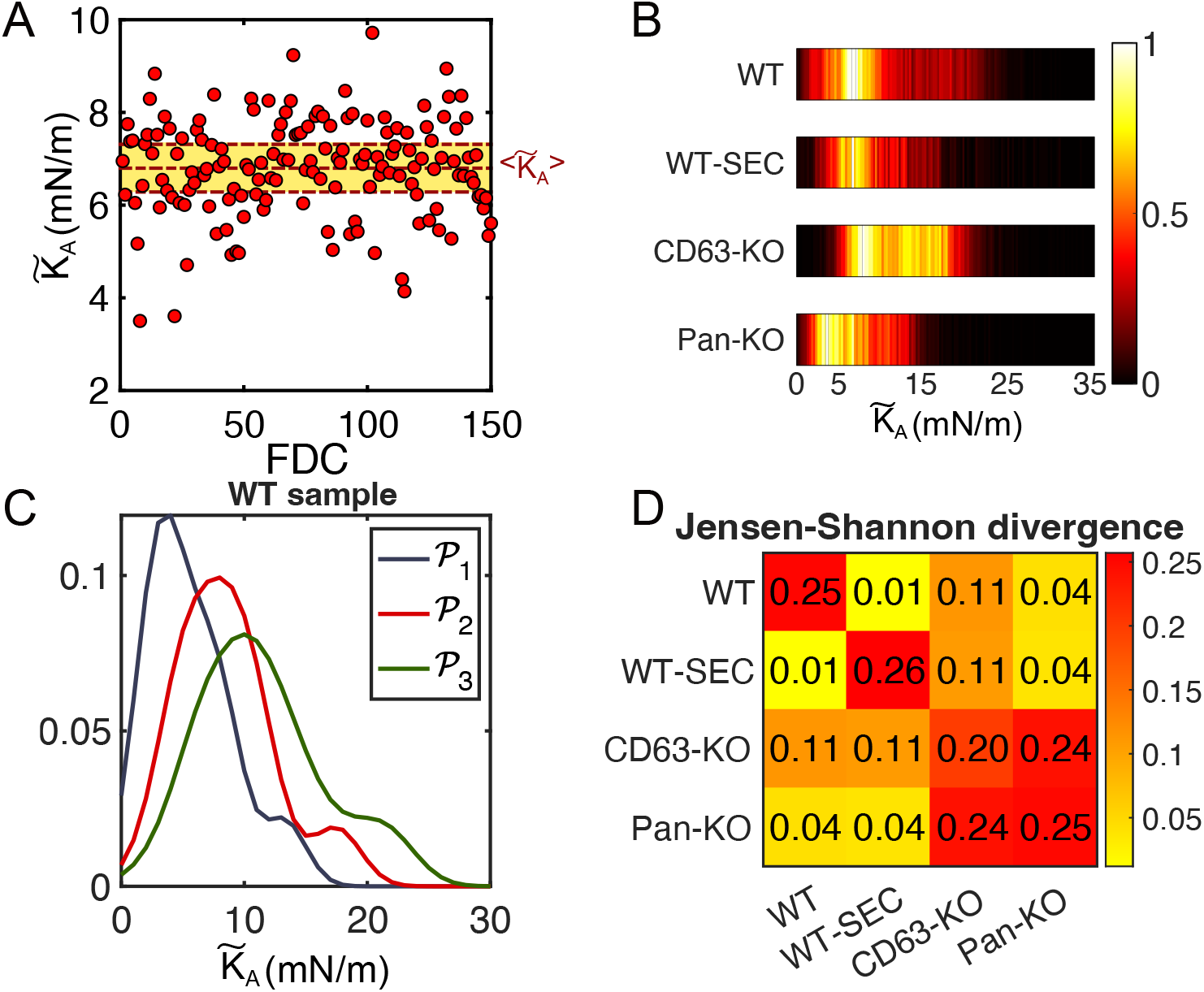
Variations of 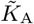 in a single vesicle. (A) The many values of 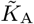 extracted from individual force–distance curve of a single vesicle. The shaded region indicates one standard deviation and the central line shows the mean value. (B) Barcode plot of all values of 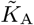 for all the vesicles in each family. (C) *𝒫*_1_, *𝒫*_2_ and *𝒫*_3_ of the WT sample. (D) Jensen-Shannon divergences between effective elastic modulus of the samples. The samples are internally compared across *𝒫*_1_ and *𝒫*_3_ (diagonal elements) while the distance across different samples (non-diagonal elements) compares *𝒫*_2_ of the two samples.

Here *D*_KL_ is the well-known Kullback–Leiber divergence. We measure the Jensen–Shanon divergence between 𝒫_1_ and 𝒫_3_ of one family and between 𝒫_2_ of two families and plot them as a matrix in Fig. (3D). The 11 element of this matrix is the *D*_JS_ between 𝒫_1_ and 𝒫_3_ for the WT family. Similarly, all the diagonal elements show the same distance for each family. The off-diagonal elements show the distance between the 𝒫_2_ of two different families. By looking at the first column (or row) of this matrix, we see that for the WT family, the distance within the family is much more than the distance between this family and any other family. There is only one case where the distance within the family is less than the distance between the families – compare CD63-KO with Pan-KO. The distance between 𝒫_1_ and 𝒫_3_ within CD63-KO family is 0.20 whereas the distance between 𝒫_2_ of CD63-KO family with Pan-KO family is 0.24. This is the only case where we may claim that our method distinguishes one family from another, even then the difference between the distances is small. Note also that we find the distance between the WT family and the WT-SEC family is very small. As reported earlier [15], additional cleaning with SEC is expected to remove protein corona. Our results mean that either the effect of the protein corona on 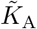is negligible or (possibly more likely) that the sample preparation steps before the AFM study, which also involves several washing steps, already removes the corona. Another aspect to consider is the compositional heterogeneity of EVs derived from a cell line or even a single cell [35, 36]. Such a variation may also result in a distribution of mechanical properties as we see in our case.

Note that we have not been able to determine *K*_A_ but an effective modulus 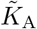. In principle, using detailed numerical simulations, as described in (Appendix B 2), it may be possible to determine *K*_A_. But, in our experience, this method is too time–consuming to apply to individual force–distance curves. We have analyzed only a fraction of the *average* force–distance curves and have obtained excellent agreement between experiment and numerical simulations. Representative force–distance curves and their fits for each family are presented in (Appendix B 2). As the number of samples measured in each family is small we have no hope of distinguishing between different families.

### D. Measurement of bending modulus

In Fig. (4A) we show a typical recording of an approach and a retraction of the AFM tip. We have already used the approach curves to find out the effective area modulus 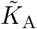. In many cases, we find that the retraction curves show the behavior seen in Fig. (4A). This indicates the formation of a tether. The retrace curve showing tether formation for GUVs [13, 37] and cells [38] usually looks different: the force remains a (negative) constant as the length of the tether increases and eventually jumps back to zero as the tether is snapped. In such cases, minimization of the free energy gives the tether force to be 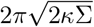 where *κ* is the bending modulus and Σ is the membrane tension. Typically, this is observed in experiments in tube–pulling assays where the GUV is aspirated in a micropipette (the aspiration pressure sets the membrane tension) while a tether is pulled. Our experimental setup is different and we observe a different qualitative behavior – the force (*F*) changes as the length of the tether (*L*) increases, *F/L* remains approximately a constant. The same behavior has already been calculated for adhered GUVs in Ref. [8]. The crucial difference between this and the tether formation in tube–pulling assays is that for the former the vesicle is not considered a quasi-infinite reservoir of lipids. This approach is appropriate for us because (i) we consider adhered vesicles and (ii) our vesicles are significantly smaller and thus, cannot be considered as a quasi–infinite reservoir of lipids. In general, the force–distance curves obtained in Ref. [8] also depends on the adhesion forces between the vesicle and the base. By assuming the volume of the tether to be much smaller than the volume of the vesicle, following Ref. [8] we obtain a simplified force– distance relation:

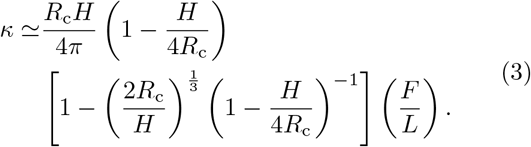

**FIG. 4.**
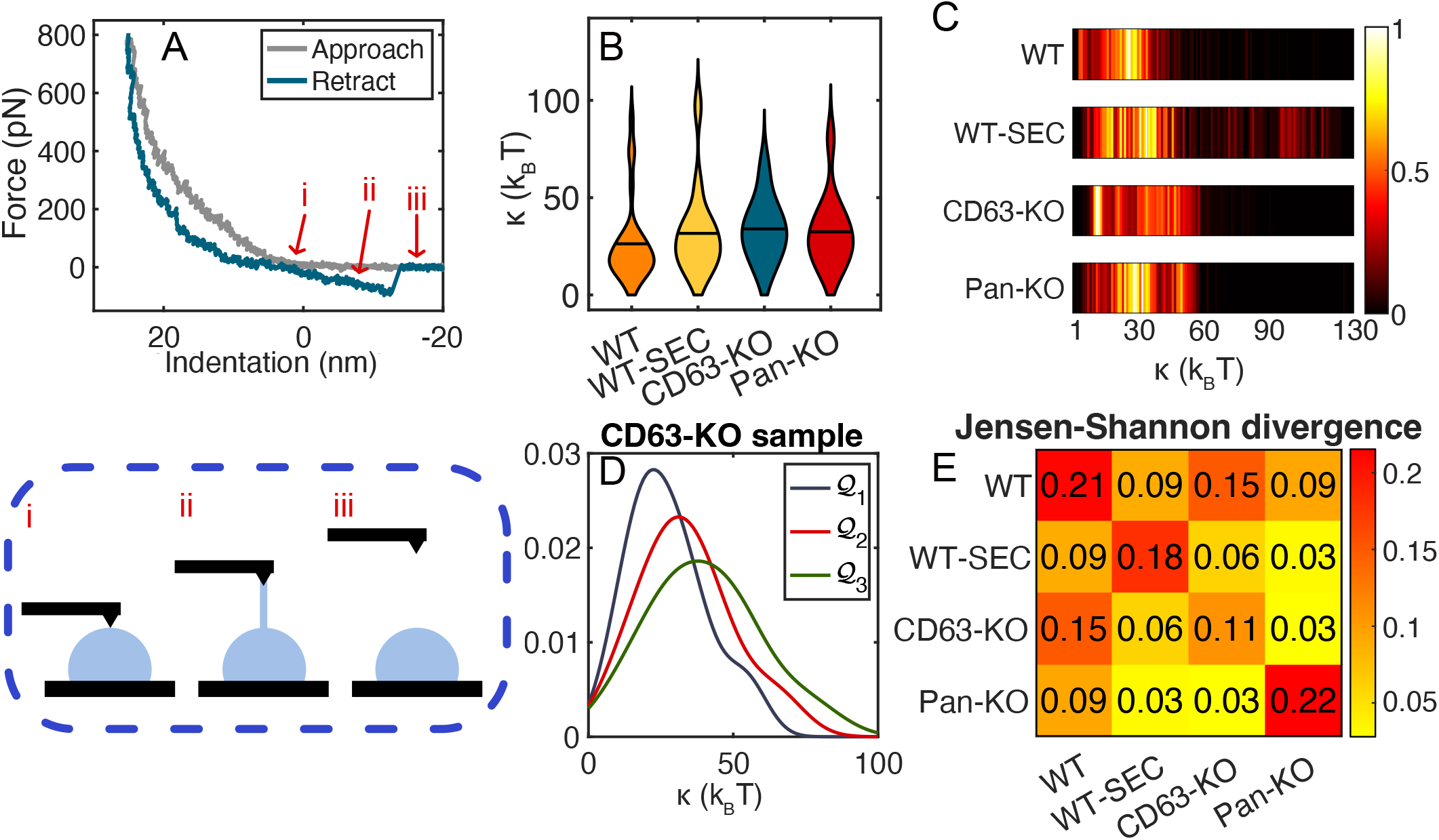
Tether formation. (A) A typical force–distance curve with the approach (grey) and retract part (blue) which shows tether formation. The different stages of tether formation correspond to the points: (i) AFM tip touches the vesicle (ii) tip is removed, tether formed, and the area of contact between the vesicle and the substrate decreases, and (iii) tether is snapped. (B) Violin plots of the bending modulus of EVs from the WT, WT-SEC, CD63-KO, and Pan-KO families. The black line indicates the mean. (C) Normalized heatmap of all extracted bending moduli for all vesicles in each sample. Bar width: 1*J/k*_B_. (D) *𝒬*_1_, *𝒬*_2_ and *𝒬*_3_ of the WT-SEC sample. (D) Jensen-Shannon divergences between effective bending modulus of the samples. The samples are internally compared across *𝒬*_1_ and *𝒬*_3_ (diagonal elements) while the distance across different samples (non-diagonal elements) compares *𝒬*_2_ of the two samples.

**FIG. 5.**
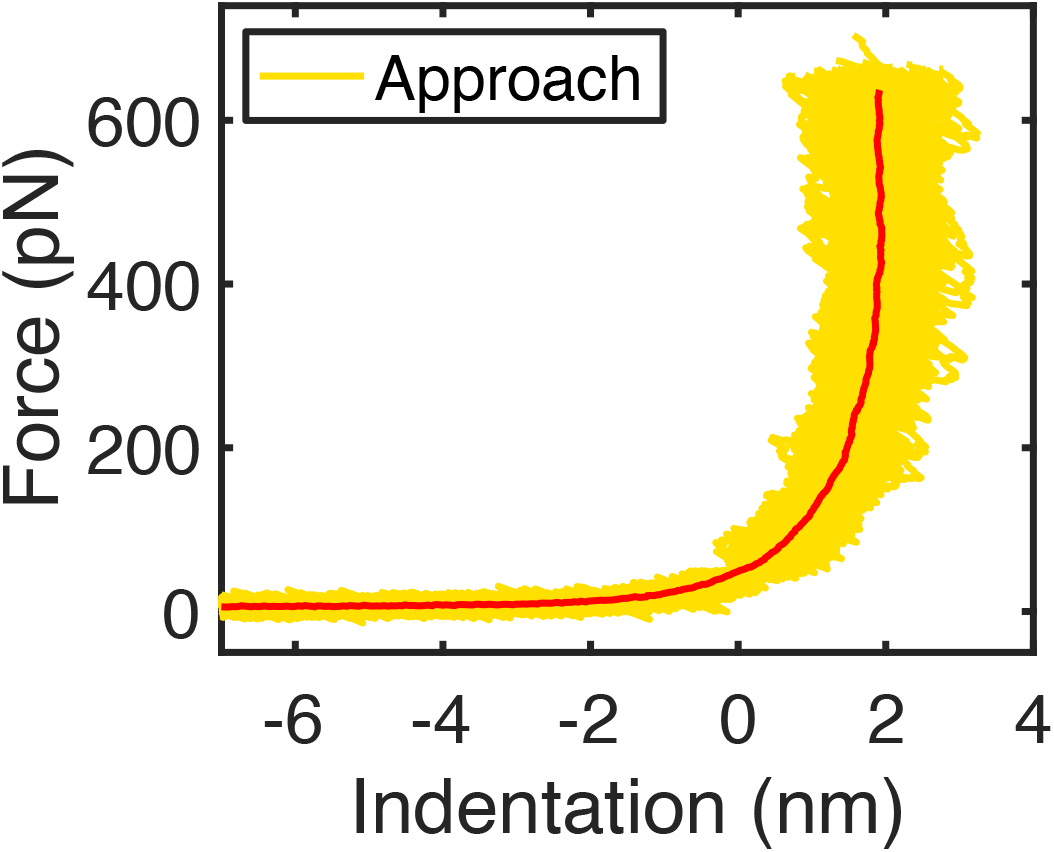
FDC on the substrate. 150 FDC superimposed on each other to show the spread of curves. The average force–distance curve is shown in red. The superposition is set so each curve has a zero indentation at 50 pN

**FIG. 6.**
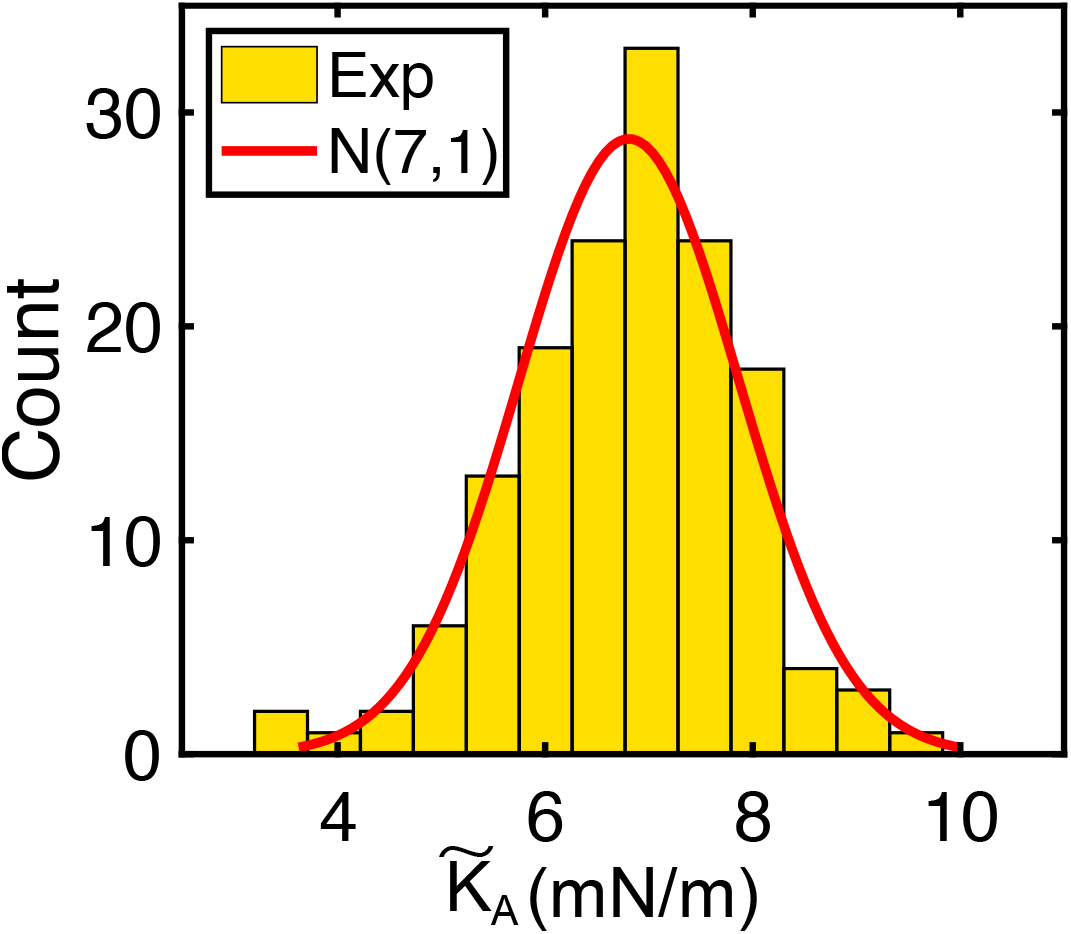
Normality of 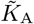 within one vesicle. Histogram plot of 150 extracted 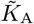 on a single vesicle. A fitted normal distribution is shown in red.

We use this expression to obtain the bending modulus *κ* from each case of the force-distance plot involving tether formation. The PDFs of the bending modulus of the families are plotted in Fig. (4B) as violin plots. In ((Appendix E)), we plot both the histograms and the PDFs estimated by the kernel density estimation for all the families. The tether formation does not happen every time the AFM tip is retracted (see discussion in IV F), therefore the sample size from which the average *κ* is calculated is smaller than in the case of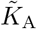. Note that, similar to what happened with the measurement of 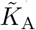, different tether formation experiments on the same vesicle yield different values for *κ*. In Fig. (4C) we show a barcode plot for all the bending moduli measured for different families. While the CD63-KO family and Pan-KO have a smaller spread in their values, the WT-SEC family shows a larger spread. Earlier measurements of the bending modulus in EVs have obtained values somewhere between 2 −20 k_B_T [11, 31, 32, 39]. Bending modulus of lipid bilayers in GUVs have been measured to be in the range of 10 − 40 k_B_T depending on composition [40–44]. The mean values of our measurements fall in the same range although, we find several values that are somewhat larger. Once again, the PDFs of our data are not single Gaussians – they also fail the Shapiro-Wilk test (Appendix C). Hence, once again, we cannot employ the t-test to check whether the differences between different families are statistically significant. We proceed in the same way as we have done for the measurement of 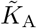. The first step is to estimate the error in the measurement of *κ*. In Eq. (3), the bending modulus depends on the height *H* and radius *R*_c_ of the vesicles and also the ratio *F/L* that we obtain from the force–distance curves. We assume that the errors in measuring *H* and *R*_c_ are small compared to the variation in *F/L* for different cases of tether formation. We attribute the source of error in *κ* to the variation in *F/L*, i.e.,

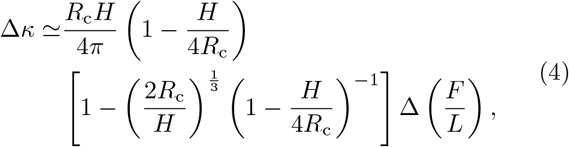

where we obtain Δ(*F/L*) from the standard deviation of *F/L* over the many measurements of tether formations. Next, we construct three PDFs for each family, one each for ⟨*κ*⟩ −Δ*κ*, ⟨*κ*⟩, and ⟨*κ*⟩ + Δ*κ*, denoted by 𝒬_1_, 𝒬_2_ and 𝒬_3_. The three PDFs for the CD63-KO family are shown in Fig. (4D). We again calculate the matrix of Jensen–Shanon distances and plot them in Fig. (4E). We find again that the diagonals of this matrix are typically larger than the off-diagonal values, i.e., the difference between the families are not statistically significant. The exception is again the family CD63-KO. The distance between 𝒫_1_ and 𝒫_3_ within CD63-KO family is 0.11 whereas the distance between 𝒫_2_ of CD63-KO family with WT family is 0.15, the PDF of bending modulus of CD63-KO is (marginally) statistically distinct from the PDF of bending modulus of the WT family.

## III. DISCUSSION

Several comments are now in order.

First concerns the theoretical model that we use to interpret the AFM measurements. We agree with earlier works [6, 11] that neither the Hertzian model nor the thin–shell–theory is appropriate to analyze the AFM measurements. We also agree that the contribution from bending is negligible. We differ from Refs. [6, 11] in one crucial way: we assume the volume of EVs to be constant, while the area changes, during the AFM indentation. We present three justifications. First, the EVs are very similar to GUVs except for their size – for GUVs the analysis of force–distance data uses the same assumption. Second, we argue that over the time scale of one force–distance measurement very little change of volume due to the osmosis of water is possible (Appendix B). And, third a straightforward dimensional argument applied to our model reproduces the linear force–distance relationship at short distance and our detailed numerical simulations produces a good fit with the experimental data. So the first take-home lesson of our work is that the large amount of already existing methods and results for GUVs are the best guide to understand the biomechanical properties of EVs with additional care necessary to take into account their small sizes and the fluctuations as explained next. Second, one crucial aspect of AFM measurement of force–distance curves of EVs that we particularly emphasize are their random variations. For a *single* EV, about 150 repeated measurements give force–distance curves that are quite different from one another, see Fig. (2A). This is a peculiar aspect of EVs – never observed in GUVs to the best of our knowledge. An important point to note is that these fluctuations seem to be Gaussian and decorrelated with one another, see Fig. (3A). We identify two mechanisms that can give rise to these fluctuations.

### 1. The thermal fluctuations

Naively, we may argue that thermal effects are not important because for lipid bilayers (*k*_B_*T/κ*) ≈1*/*20 is less than unity. Recent theoretical works [10, 45, 46] have demonstrated that in thin shells the dimensionless number that determines the importance of thermal effects is the elasto-thermal number, given by 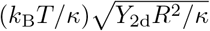 where *Y*_2d_ the *two-dimensional* Young’s modulus. These results cannot be directly applied to EVs because EVs have a fluid membrane whereas the thin–shell theory considers a solid membrane. Nevertheless, for EVs, we can define a similar elasto-thermal number by replacing *Y*_2d_ by 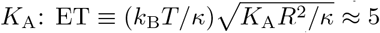, where we have used [41] *K*_A_ ≈ 200 mN m^−1^ and *κ* ≈ 10^−19^J, and *R* = 100 nm. The elasto-thermal number is larger than unity, hence we do expect thermal effects to be important.

### 2. The diffusion of membrane molecules

It is well known that membrane proteins diffuse on the lipid membrane [47, 48]. The diffusion constant, *D*, is estimated to vary over a large range [49, Table 1]. Let us take a representative value [50] of *D* ≈ 1 μm^2^ s^−1^. The typical time scale of a complete force–distance measurement is about 1 s. Over that time scale a protein diffuses approximately over an area of 1 μm^2^, which is significantly larger than the surface area of the EVs. This implies that by a single force–distance measurement, we obtain some kind of average elastic modulus. Thus every force–distance measurement yields a slightly different value of elastic modulus. However, the estimate of *D* we have used might be inaccurate because: (i) proteins on the surface of EVs are known to form clusters [51]; (ii) the value of *D* has been measured for almost flat surfaces [49], for example on surfaces of cells; (iii) in addition, the interaction between proteins with the AFM tip and the surface may affect their diffusion. Note also that if the characteristic time scale of diffusion of proteins slowed down so much that it became comparable to the typical time scale of a force–distance measurement, then we would expect some correlation between successive measurements. But we do not see this.

**TABLE I.** The *K*_A_ and Σ_0_ values for four families extracted using our numerical model.

| Family | EV | $K_A$ (mN/m) | $\Sigma_0$ (mN/m) | Family | EV | $K_A$ (mN/m) | $\Sigma_0$ (mN/m) |
| --- | --- | --- | --- | --- | --- | --- | --- |
| WT | 221216ev1 | 299.83 | 10.20 | WT-SEC | 230619ev11c | 799.64 | 12.07 |
|  | 221216ev6 | 424.32 | 7.28 |  | 230619ev3 | 213.08 | 14.08 |
|  | 221222ev2 | 1513.60 | 4.35 |  | 230622ev4 | 645.21 | 11.01 |
|  | 221222ev8 | 303.97 | 8.60 |  | 230628ev2 | 1170.42 | 11.57 |
|  | 230718ev5 | 1631.79 | 11.54 |  | 230717ev6 | 300.39 | 33.47 |
| CD63KO | 221219ev2 | 391.87 | 7.40 | Pan-KO | 221213ev4 | 332.88 | 5.27 |
|  | 221219ev3 | 960.15 | 7.60 |  | 221213ev7 | 648.60 | 8.58 |
|  | 221219ev7 | 673.43 | 17.38 |  | 230118ev3 | 695.67 | 0.80 |
|  | 221219ev9 | 240.20 | 17.00 |  | 230118ev4 | 474.98 | 2.18 |
|  | 230130ev11 | 451.35 | 8.89 |  | 230118ev7 | 187.31 | 1.46 |

At the moment, it is not clear to us, which of these two mechanisms makes the dominant contribution. During the last couple of years, there have been several measurements of the nanomechanical properties of vesicles through atomic force microscopy [6, 30, 39, 52–55]. Most of these rest on the assumption that one or few measurements on each vesicle is enough for a representative value. Here we show explicitly and justify theoretically that this is not the case.

Note that another consequence of such protein diffusion is that if antibodies are used to capture the EVs, all the complementary proteins may eventually diffuse to the substrate side, thus inducing some degree of bias to the compositional fluctuations. To reduce and possibly to eliminate such a scenario, we use electrostatics capture of EVs, as we described in our previous work [56]. This approach makes the selection of EVs less biased.

The next remark is about tether formation. Typical theory of force–distance curves for tether formation in GUVs [37] and cells [38] shows that the force remains a constant as the length of the tether increases and eventually jumps back to zero as the tether is snapped. Typically, this is observed in experiments in tube–pulling assays where the surface tension of the membrane is held constant. Nevertheless, the same theory has been used to interpret tether formation of adhered cells [57] and vesicles [11, 13, 19, 31] in which case the surface tension cannot be controlled. In our experiments neither can we control the surface tension – a direct consequence of adhering the vesicles – nor can we assume that the EVs have a *reservoir of lipids* due to their small sizes. We obtain a linear force–distance relationship. We follow Ref [8], whose theory gives a linear force–distance relationship which we use to measure the bending modulus. To the best of our knowledge, there has been only one experimental measurement [9] of tether formation in GUVs that found a linear force–distance relationship although Ref. [9] suggested a theoretical interpretation different from Ref. [8]. The linear force–distance in tether formation is also found in bacterium [58], and fibroblasts [59], but in these cases the mechanism may be different. To summarize, ours is one of the first experimental observation of linear force–distance relationship which may be interpreted using the theory of Ref. [8].

Note also the natural heterogeneity of biomechanical properties within a family of EVs. Our data show that the PDF of the elastic modulus is not likely to be Gaussian but can be best represented by a superposition of several Gaussians. From the biological point of view, this is expected, because EVs are known to be heterogeneous containing several different subpopulations, each of which may generate a Gaussian distribution of biomechanical properties. This absence of simple Gaussianity also demands statistical methods that are suitable for families of EVs. As the underlying normality assumption for the t-test is violated we use the concept of Jensen-Shannon divergences to differentiate between probability distributions. We believe this is better suited as a statistical approach due to the natural variations of the elastic moduli of EVs.

Now let us discuss to what extent our original intention, v.i.z., to differentiate EV subpopulations based on their elastic properties, has been served. Membrane protein-based discrimination of EV subpopulation is by far the most investigated topic in the field, thus also providing the basis for our primary question, v.i.z., does the deformability of EVs depend on their membrane protein composition? We investigate this question by knocking out three tetraspanins which are highly abundant in HEK293T-derived EVs [60], thus they become an obvious choice for our experimental design. Note that knocking out of certain proteins may also have unwanted consequences where the proteins are replaced by other proteins or lipids. For example, it may change the cholesterol content of the EVs [61], which in turn can change their elastic properties, [see, e.g., 62, 63]. Furthermore, Ref. [64] has suggested that the presence of proteins may change the local curvature of the lipid membrane thereby changing its elastic properties [64]. Our results suggest that the natural variation in the elastic moduli within a family is too large to detect changes, induced by protein alteration, between families. The only exception is the CD63-KO family, which shows a statistically significant difference. The final take-home message of our work is that the error in earlier experiments may have been underestimated.

The study of the nanomechanical properties of extracellular vesicles is still in the early stages. The central goal of our paper is to address one of its greatest challenges: to properly identify stable and relevant protocols of analysis of EVs.

## IV. METHODS

### A. Chemicals and materials

High-purity deionized water (DIW) with a resistivity of 18 MΩ cm was used throughout all the experiments. Casein (C5890) in powder, phosphate-buffered saline (PBS; P4417) in tablets, and poly-l-lysine (PLL; P2636) were purchased from Sigma-Aldrich (Burlington, MA, United States).

### B. Cell culture and extraction, purification, and isolation of EVs

HEK293T (human embryonic kidney-293T) cells were propagated in Dulbecco’s modified Eagle’s medium (DMEM) containing Glutamax-I and sodium pyruvate; 4.5 g*/*L Glucose; Invitrogen) supplemented with 10% fetal bovine serum (FBS; Invitrogen) and 1% Antibiotic-Antimycotic (Anti-Anti; ThermoFisher Scientific). 48 h prior to collection of conditioned media for EV isolation from HEK293T cells, cells were washed with PBS and the medium was changed to OptiMem (Invitrogen) [65]. HEK293T-CD63KO and HEK293T-PAN-KO (CD9, CD63, and CD81) cell lines were generated by the delivery of Cas9-gRNA ribonucleoproteins (IDT, Integrated DNA Technologies, Coralville) targeting respective tetraspanin sequences using RNAiMAX (Thermo Fisher Scientific, Waltham, MA, USA). Shortly, 10,000 cells per well were seeded in a 96-well plate and treated after 24 h with 100 ng of Cas9-RNAiMax per well, following the protocol of Chesnut et al 2015[66]. Three days after treatment the cells were stained with anti-CD63-APC or a mixture of anti-CD9/63/81-APC antibodies [14], and successfully edited cells were sorted on a BD Fusion flow cytometric cell sorter as single cells per well into 96-well plates. The resulting colonies were validated and expanded as stable cell lines. All cell lines were grown at 37°C, 5% CO2 in a humidified atmosphere and regularly tested for the presence of mycoplasma. For EV preparation, cell culture-derived conditioned media (CM) was first pre-cleared from cells and debris by low-speed centrifugation (700 × *g* for 5 min) and subsequent centrifugation at 2,000 × *g* for 20 min to remove larger particles and debris. Next, media was filtered through 0.22 μm bottle top vacuum filters (Corning, cellulose acetate, low protein binding) to remove any larger particles. Precleared CM was subsequently concentrated via tangential flow filtration (TFF) by using the KR2i TFF system (SpectrumLabs) equipped with modified polyethersulfone (mPES) hollow fiber filters with 300 kDa membrane pore size (MidiKros, 370 cm^2^ surface area, SpectrumLabs), at a flow rate of 100 mL/min (transmembrane pressure at 3.0 psi and shear rate at 3700 sec^−1^, as described previously [67]. WT-SEC EVs were additionally purified by bind-elute size exclusion chromatography (BE-SEC): CM were concentrated by TFF as described above and then loaded onto BE-SEC columns (HiScreen Capto Core 700 column, GE Healthcare Life Sciences), connected to an ÄKTAstart chromatography system (GE Healthcare Life Sciences) as described previously [67]. Amicon Ultra-0.5 10 kDa MWCO spin-filters (Millipore) were used to concentrate the sample to a final volume of 100 μL. EVs were stored in Maxymum Recovery polypropylene 1.5 mL tubes (Axygen Maxymum Recovery, Corning, cat MCT-150-L-C) in PBS-HAT buffer (PBS, 25 mM Trehalose, 25 mM HEPES, 0.2% Human Serum Albumin) before usage as described previously [14]. Prepared sEV samples were characterized by Nanoparticle tracking analysis (NTA) to determine particle size and concentration using the NanoSight NS500 instrument equipped with NTA 2.3 analytical software and an additional 488 nm laser [67]. The samples were diluted in 0.22 μm filtered PBS to an appropriate concentration before being analyzed. At least five 30-second videos were recorded per sample in light scatter mode with a camera level of 11-13. Software settings were kept constant for all EV measurements (screen gain 10, detection threshold 7). The analysis was performed with the screen gain at 10 and the detection threshold at 7 for all EV measurements. Successful knockout of CD63 and CD9/CD63/CD81, respectively, were validated by multiplex bead-based EV flow cytometry [60] and single EV imaging flow cytometry as described before [68] (data not shown).

### C. EV sample preparation

After purification and isolation, the EVs were adhered to coverslips. The coverslips were cleaned from organic residue with a 5:1:1 RCA-1 solution of deionized water, NH_3_, and H_2_O_2_ (90°C, 10 minutes). After cleaning, the coverslips were coated with 0.001% poly-l-lysine for 90 seconds. 100 μL of 1 × 10^9^ part*/*mL of the sample suspended in 1× PBS was then incubated for 1 hour at room temperature. A cut-up four-well insert (80,469, ibidi GmbH, Gräfelfing, BY, DE) was used to limit the spatial spread of the EV sample on the coverslip. After incubation, the substrate was thoroughly washed with 1× PBS. To reduce the risk of unsolved salt crystals contaminating scans, the PBS was filtered through a 0.2 μm PTFE filter (514-0070; VWR; AB, SE). Before loading the sample on the AFM, the liquid volume was increased to approximately 400 μL to avoid drying out the sample and deflating the vesicles during imaging. The dried-up sample showed up either as great spikes in the topographical image or as elongated, short vesicles (Appendix F).

### D. Atomic Force Microscopy (AFM)

The biophysical measurements were performed with a NanoWizard 3 BioScience AFM from JPK (Berlin, BE, Germany). A CoverslipHolder from Bruker was used to enable a liquid environment. Performing measurements in 1 × PBS allows a higher preservation of their spherical shape than measurements performed in air [69]. Qp-BioAC cantilevers from NanoAndMore GMBH (Wetzlar, HE, Germany) with nominal spring constants of 0.06 N*/*m, 0.1 N*/*m, and 0.1 N*/*m were used. All sessions started using the cantilever with the lowest nominal spring constant which was swapped to the other cantilever as tip contamination occurred. Before imaging the spring constant of the cantilever was determined through the thermal noise method [70]. Images were obtained in Quantitative Image (QI) mode with an imaging force of 200 pN. First, a coarse scan of a 10 μm ×10 μm large region with a resolution of 39 nm/pixel was carried out. Objects with height greater than 25 nm and width greater than 100 nm at this resolution were classified as potential EVs. For this, a 500 nm ×500 nm or 400 nm × 400 nm scan centered around the object at a resolution of 3.9 or 4 nm pixel^−1^ was considered. If the object exhibited a spherical cap shape it was assumed to be a vesicle, and force spectroscopy measurements were performed at its center. The force spectroscopy measurements were carried out with the maximum indentation force of 0.8 nN. At least 150 consecutive indentations were performed at the constant speed of 1 μm*/*s. After the set of force spectroscopy measurements, the vesicle was imaged again to ensure that the measurements had not damaged, deformed, or otherwise moved the vesicle.

### E. Post data capture processing

In Gwyddion, all single EV images were tilt-corrected and shifted in the z-direction to let the lowest pixel represent zero level. A line profile in the slow scan direction over the EV peak was exported to Matlab where a script calculated a radius of curvature (*R*_c_) by fitting a circle to all the points with heights greater than half the maximum height as previously suggested in Ref. [19]. In the same protocol, Vorselen et al., suggest applying forces in the 5-10 nN range to completely indent the vesicle and reach the substrate. This allows the height, *H*, of the vesicle to be extracted from the force-distance curve instead of through the topographical images. However, such large forces are likely to rupture the vesicles and we avoid this step. Instead, the height was extracted from the line profile over the center of the EV.

### F. Force distance curve analysis

The force-distance curves were processed in the JPK Data Processing program. First, the offset and the tilt of the baseline were corrected. After this, the contact point between the AFM tip and the vesicle was found. Finally, the height was corrected for cantilever bending. A successfully processed curve would have its contact point at 0 nm indentation, and a constant baseline around 0 pN. The stiffness was extracted as the slope of the approach curve in the penetration region 0.05 − 0.1H. The retract curves were exported to Matlab where a script identified curves with tether formations and extracted their length and force magnitude. A curve would be ascribed with a tether if the global force minimum occurs at *<* 0 nm indentation, has a greater magnitude than 100 pN, and the force returns to the baseline before the AFM tip is returned to its original state. The set of selection rules for the force–distance curves are given in (Appendix G).

### G. Statistical analysis

Statistical significance was established using an unpaired, two-sample t-test at *α* = 0.05. Probability density functions were generated and compared with the Jensen-Shannon distance. All the statistical tests were scripted using MATLAB^®^, except for the Shapiro-Wilk goodness-of-fit test which was performed in R.

## ACKNOWLEDGMENTS

FS, HK, PM, and AD acknowledge the financial support of the Erling-Persson Family foundation and Stockholm County Council through grant FoUI-966345. DM, VP, and VA acknowledge the financial support of the Swedish Research Council through grants 638-2013-9243 and 2016-05225. DM sincerely thanks Sreekanth Manikandan for the discussions. NORDITA is partially supported by NordForsk. AG is an International Society for Advancement of Cytometry (ISAC) Marylou Ingram Scholar 2019-2024 and supported by a Karolinska Institutet Network Medicine Alliance Collaborative Grant.

## Appendix A Force distance curves (FDC)

## Appendix B Theory of elastic deformation of EVs

### 1. Theory for small deformation

Let us start with a vesicle adhered to a glass plate. We use a set of Cartesian coordinates *X*_1_, *X*_2_, *X*_3_ such that the plate is given by plane *X*_3_ = 0. We also introduce a generalized coordinate *x*_1_ and *x*_2_ the describe the surface of the vesicle [see e.g., 71]. We start with the configuration of the adhered EV on the glass surface given by the function

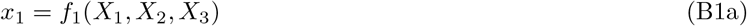

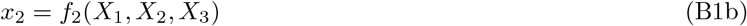

where the functions *f*_1_ and *f*_2_ can be inverted to obtain

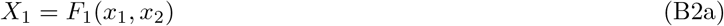

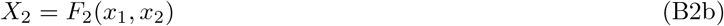

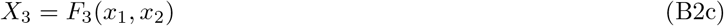

For example, if the undeformed shape of the vesicle where a perfect sphere just touching the glass plate, then we could use *X*_1_ = *x, X*_2_ = *y, X*_3_ = *z* where *x, y, z* are the usual Cartesian coordinates with *z* = 0 the coordinate of the glass plate. We could use as generalized coordinates the usual angles on the surface of the sphere: *x*_1_ = *θ* and *x*_2_ = *φ* such that

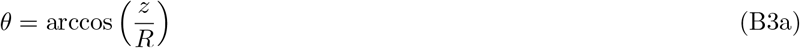

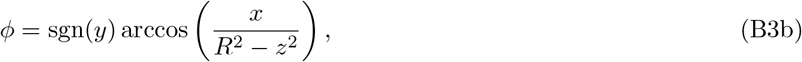

where 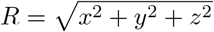 is the radius of the sphere. These equations can be inverted to obtain

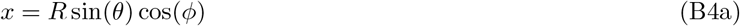

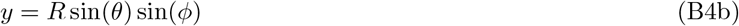

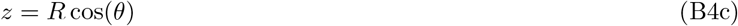

The deformation of the membrane relative to this configuration can be described by an in-plane (two-dimensional) vector ***u***(*x*_1_, *x*_2_) and an out-of-plane (scalar) deformation *h*(*x*_1_, *x*_2_). The free energy of the membrane is given by [22]

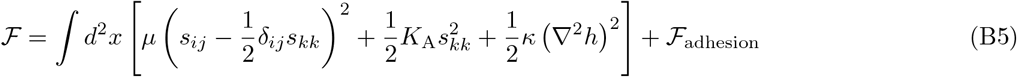

Where *μ* is the in-plane shear modulus and *K*_A_ is the in-plane area modulus and ℱ_adhesion_ is the free energy contribution from adhesion. The area element 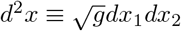 where *g* is the determinant of the metric tensor associated with the metric for the adhered shape. The strain tensor *s*_*ij*_ is given by

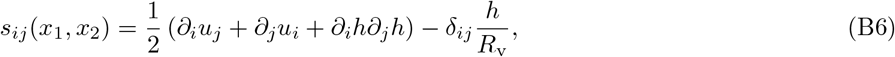

where *R*_v_, an effective radius of the vesicle can be set by 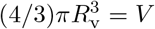 the volume of the vesicle which is assumed to be constant. As we consider a fluid membrane we set *μ* = 0 in (1). Let us now consider the AFM tip applying a force *F* that causes a deformation *d*. The deformation is very small compared to the *R*_v_ such that *d/R*_v_ is a small parameter. The change in free energy can be estimated as

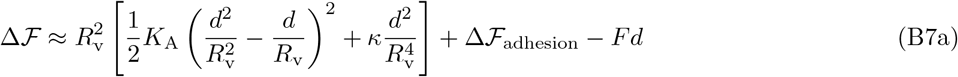

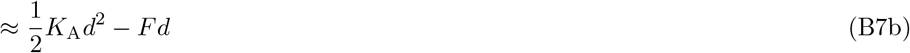

In (B7a) we assume that the gradients of the in-plane deformation to be very small and estimate the typical value of out-of-plane deformation *h* by *d* and the the typical derivatives of *h* by *d/R*_v_ and the integral over the surface of the vesicle as 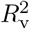. The last term in (B7a) is the work done on the vesicle by the AFM. It is negative because the force acts inward. To arrive at (B7b) from (B7b) we :

a. Expand the parenthesis in (B7a) and keep only the leading order contribution in *d/R*_v_
b. Ignore the change in free energy of adhesion for small *d*
c. Ignore the contribution from bending as it is a higher-order in the small parameter

Equation (B7b) is correct up to coefficients that depend on the geometry of the indenter and the adhered shape of the vesicle. Minimizing Δ ℱ with respect to *d* we obtain *F* ∝ *K*_A_*d*. Although this gives a linear force–distance relationship for small deformation this does not allow us to determine the elastic constant *K*_A_ because the constant of proportionality, which depends on the adhered shape of the vesicle and the geometry of the indenter, can not be determined by the kind of dimensional arguments given here. We need detailed numerical simulation.

#### 2. Numerical simulation

In this subsection, we present our simulations of a vesicle on a substrate indented by a conical indenter to calculate the force–distance curve. This section is added for completeness. We closely follow Ref. [29, 72]. The calculation operates under the following assumptions:

1. The pressure difference across the membrane is constant.
2. The total volume is conserved, see section B 3 for justification.
3. The deformation is axially symmetric.
4. The membrane tension is uniform.
5. The bending rigidity is negligible.

The model is shown in Fig. (7). We divide the shape into 4 regions, ℛ_i_ ≡ (*s*_i_ → *s*_i+1_), where i goes from 1 to 4. The region R_0_ is the contact region with the flat substrate. We use Young-Laplace law:

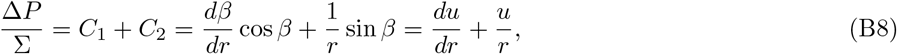

where Δ*P* is the pressure difference across the membrane, Σ is the tension, *C*_1_, *C*_2_ are the principal curvatures and *u* ≡ sin *β*. Since Δ*P/*Σ is constant, we can solve Eq. (B8) for each region to get:

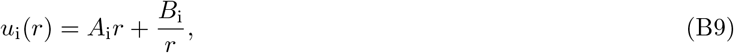

where *A*_i_, *B*_i_ are the constant of integration for i-th region. We implement the following boundary conditions:

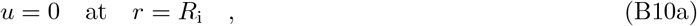

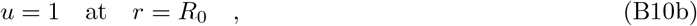

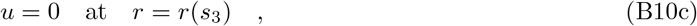

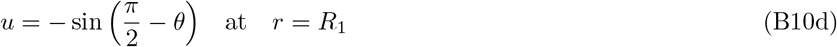

to obtain

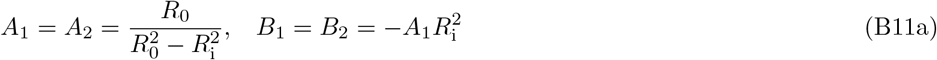

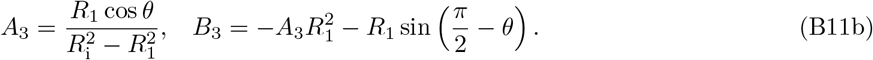

**FIG. 7.**
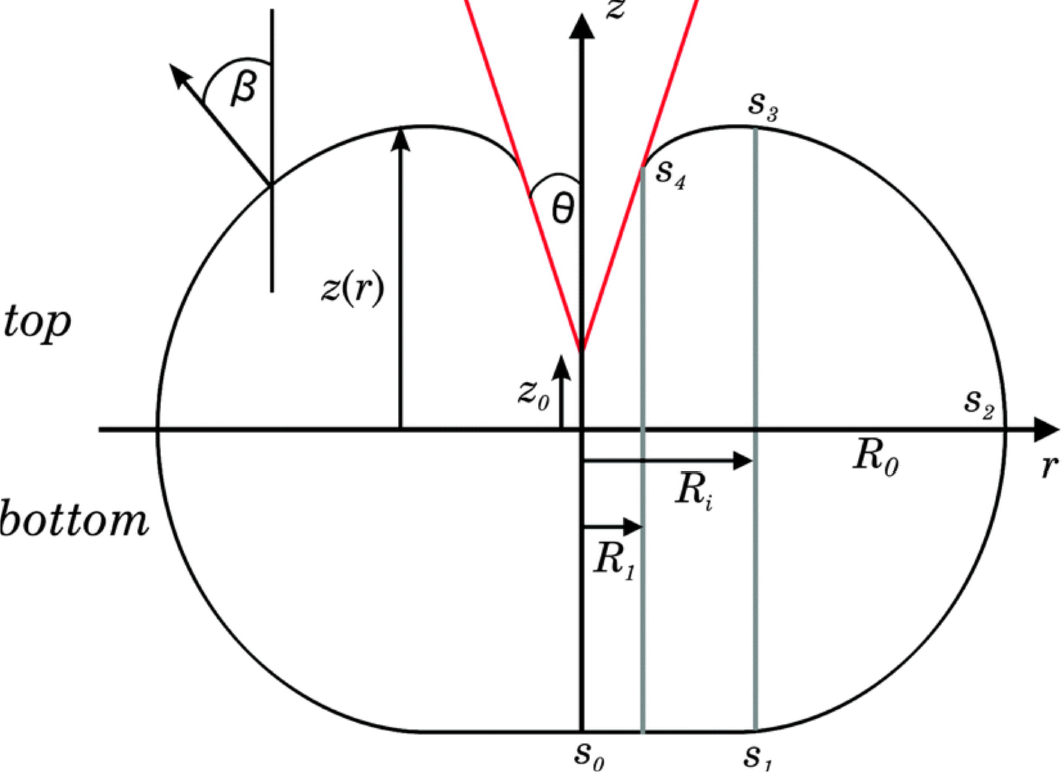
Schematic of a vesicle on a substrate indented by a conical indenter. Figure reprinted with permission from [29].

Right now we have three variables; *R*_0_, *R*_i_, and *R*_1_, but we have two additional constraints as well. First, the force from the indenter *F*_top_ and the adhering force *F*_bottom_ should be equal. We write

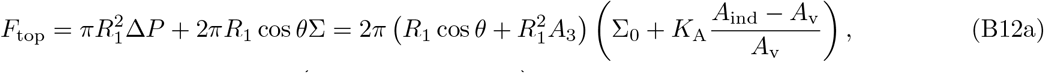

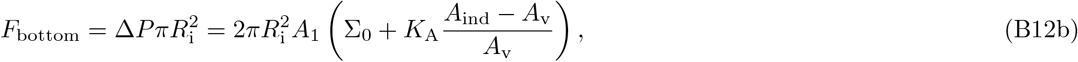

where Σ_0_ is the pre-tension, i.e., tension before indenting. This is an effective parameter to take into account the fact that before indenting the vesicle is adhered to the surface and not a perfect sphere. The area before indentation is *A*_v_ and

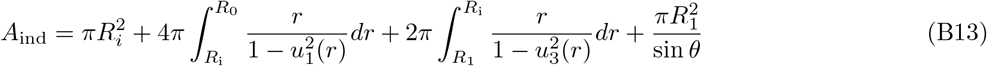

is the area of the vesicle after indentation. By setting *F*_top_ = *F*_bottom_, we get

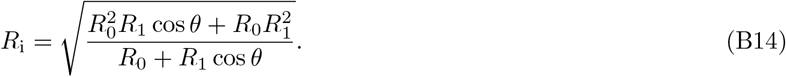

Additionally, the volume of the vesicle

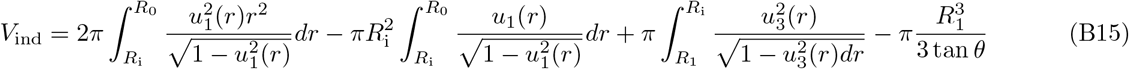

is also conserved. We satisfy this constraint numerically. The indentation depth, d, in terms of *R*_0_, *R*_i_ and *R*_1_ is

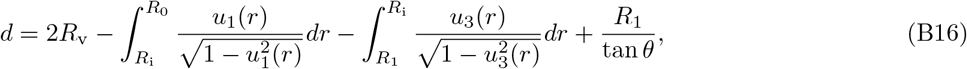

where *R*_v_ is the radius of the vesicle before deformation. We start with some values of *R*_0_, and use constraint in (B15) and (B14) to compute the vesicle shapes for different indentations. We compute *d* and *F* using (B16) and (B12b) respectively. Thus we get a force-distance curve for a given *K*_A_ and Σ_0_.

The procedure to deduce the values of *K*_A_ and Σ_0_ from a given experimental curve is as follows:

1. We shift the experimental data in the horizontal direction such that the zero of the force-distance curve coincides with the origin of the coordinate system. We find the zero by fitting a straight line to the first 10% of the data. We call the experimental force–distance curve ***F*** ^exp^.
2. We start with a guess for *K*_A_ and Σ_0_, (*K*_A_, Σ_0_) = (0.5, 0.005) (N/m), and solve our numerical model to obtain a force–distance curve which we call ***F*** ^sim^.
3. We then calculate the error 𝔼 = ||log(***F*** ^exp^) − log(***F*** ^sim^) ||, where ***F*** ^exp^ and ***F*** ^sim^ are the experimental and simulated force values respectively and ||· || represents the second-norm. Note that, we use the logarithm of the force in error computation because the force values range over three decades and we want a balanced representation of both the linear (small forces) and nonlinear (larger forces) regimes.
4. We use the `curve_fit` module of the scipy library and employ the Levenberg–Marquardt (LM) optimization scheme [73, 74] to arrive at a pair of values of *K*_A_ and Σ_0_ that minimize the error above.
5. We reject the values of *K*_A_ and Σ_0_ if: (a) the error is more than 2% of the experimental force values or (b) the optimized *K*_A_ and Σ_0_ values are negative. Note that the optimization problem is not convex. There is no guarantee that there is one unique solution.

In our experience, this method of analysis cannot be automated, every case has to be considered separately. Also note that we do not know the numerically obtained force–distance curve as a function that can be evaluated at every step of the optimization process. We must solve our numerical model for many values of the imposed force at every step of the optimization process. As this is very time-consuming it is impossible to use this technique to extract *K*_A_ and Σ_0_ from individual force–distance curves and construct a PDF as we have done for 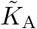. We have analyzed only a fraction of the *average* force–distance curves and have obtained excellent agreement, presented in Fig. (8). We show in Table B 2 the values of *K*_A_ and Σ_0_ we obtain.

**FIG. 8.**
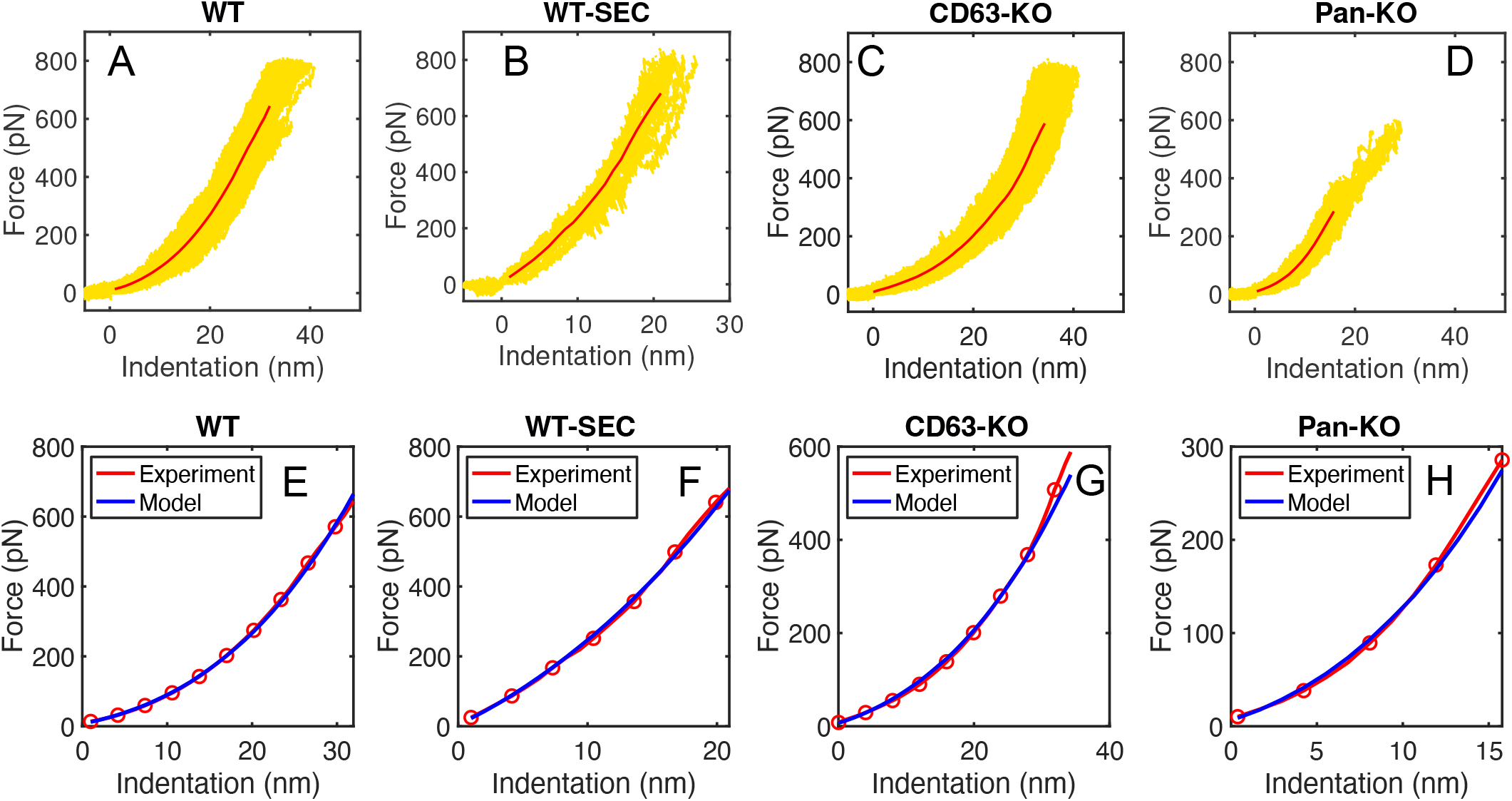
Force distance curves (yellow) and their average (red) for a representative vesicle from the (A) WT family, (B) WT-SEC, (C) CD63KO, and (D) Pan-KO. (E-H) The average experimental force–distance curve (red) and its numerical fit (blue).

**FIG. 9.**
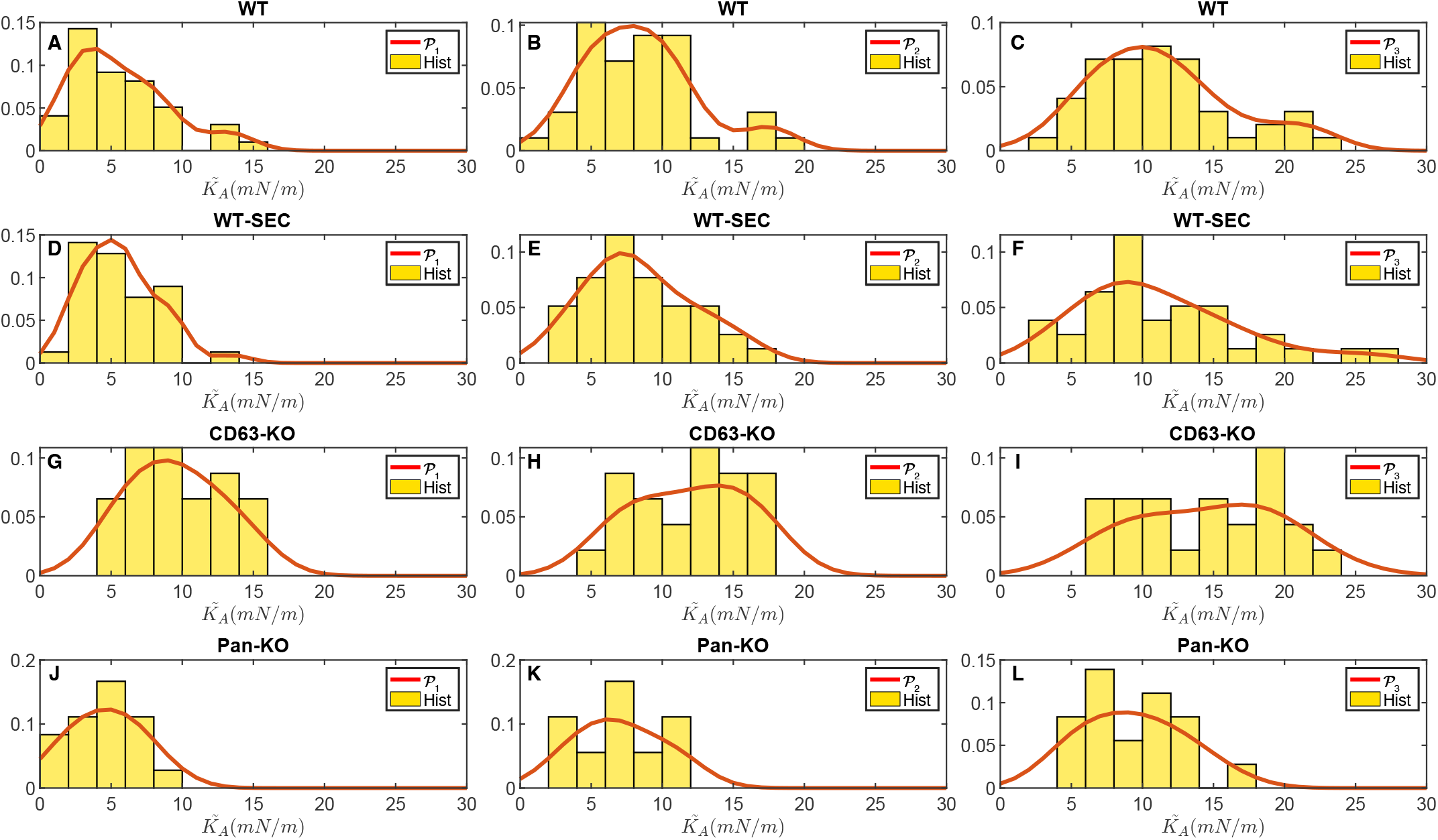
Effective elastic modulus. Histogram of linear stiffnesses and their corresponding probability density function. Each row corresponds to a specific sample and each column a way to achieve the linear stiffness values. (A-C) show the histogram and PDF for the WT sample, (D-F) WT-SEC, (G-I) CD63-KO, and (J-K) for Pan-KO. Probability density functions generated with kernel smoothing function estimate.

**FIG. 10.**
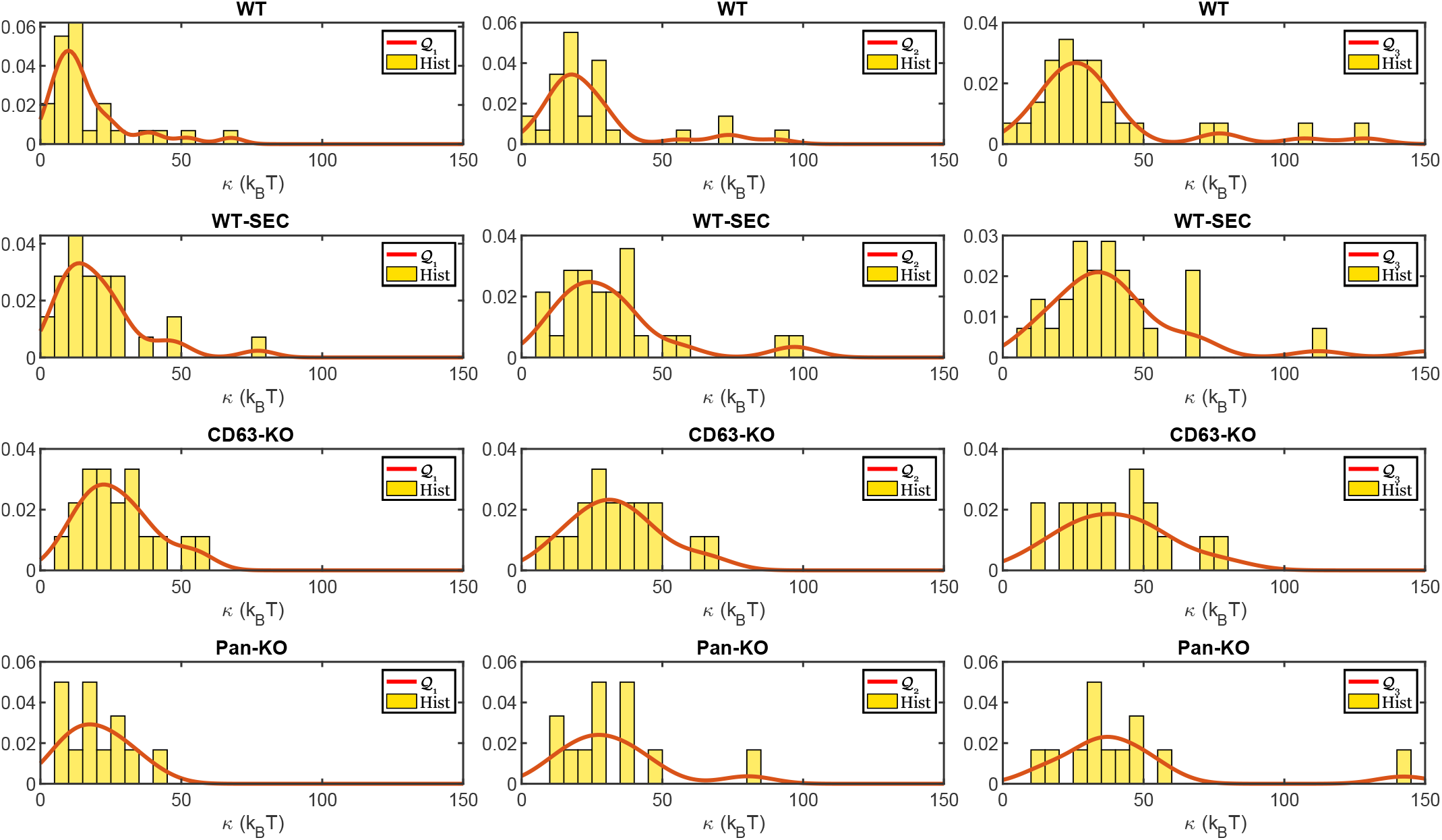
Bending modulus. Histogram of bending modulus and their corresponding probability density function. Each row corresponds to a specific sample and each column is a way to achieve the linear stiffness values. (A-C) show the histogram and PDF for the WT sample, (D-F) WT-SEC, (G-I) CD63-KO, and (J-K) for Pan-KO. Probability density functions generated with kernel smoothing function estimate with positive support and bandwidth = 0.5.

**FIG. 11.**
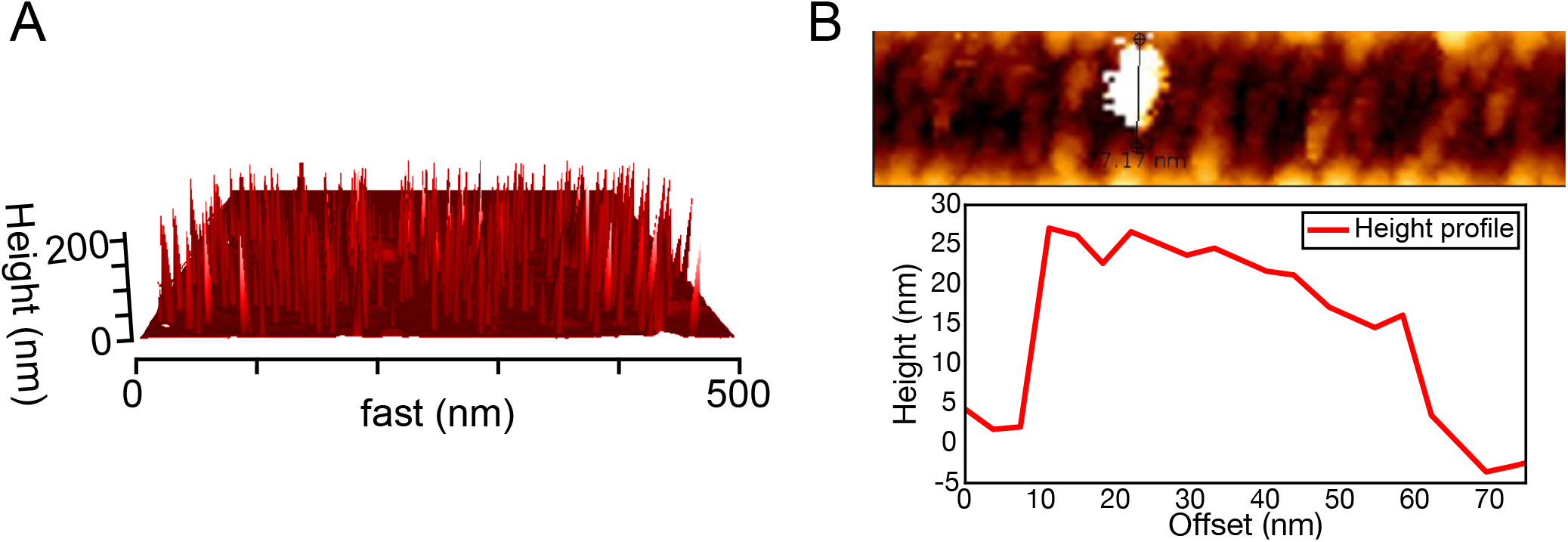
Problems caused by dried sample. Topographical images showing (A) small height spikes and (B) a vesicle with a deflated height profile.

For synthetically made GUVs the *K*_A_ has been measured using micropipete aspiration. The measured values range from approximately 150 mN/m to 2000 mN/m depending on their composition [75, Table 11.1 and 11.2]. For GUVs composed of DOPC lipids Ref. [29], whose method we use, obtain an average value of 40 mN/m. The values we obtain are approximately in the same range. As the number of samples measured in each family is too small we have no hope of distinguishing between different families in this manner.

**TABLE II.** Shapiro-Wilk goodness-of-fit test. Resulting p-values from the Shapiro-Wilk test for the different samples and the two calculated mechanical properties. H_0_ = The data is from a normal distribution.

| P-value | WT | WT-SEC | CD63-KO | Pan-KO |
| --- | --- | --- | --- | --- |
| Linear stiffness | 0.00 | 0.24 | 0.17 | 0.36 |
| Bending modulus | 0.00 | 0.00 | 0.80 | 0.07 |

**TABLE III.** Bandwidth. Bandwidth of the generated PDFs in Fig. 9. Selected with Silverman’s rule of thumb.

| Bandwidth | $\mathcal{P}_1$ | $\mathcal{P}_2$ | $\mathcal{P}_3$ |
| --- | --- | --- | --- |
| WT | 1.56 | 1.89 | 2.27 |
| WT-SEC | 1.24 | 1.98 | 2.63 |
| CD63-KO | 1.80 | 2.12 | 2.33 |
| Pan-KO | 2.24 | 2.55 | 3.41 |

**TABLE IV.** Area under curve. Area under the 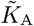 probability density functions from the kernel density estimator. The value of 1 indicates a proper probability density function.

| | $\tilde{K}_A - \Delta\tilde{K}_A$ | $\tilde{K}_A$ | $\tilde{K}_A + \Delta\tilde{K}_A$ |
| --- | --- | --- | --- |
| WT | 0.9714 | 0.9926 | 0.9952 |
| WT-SEC | 0.9922 | 0.9906 | 0.9854 |
| CD63-KO | 0.9977 | 0.9987 | 0.9952 |
| Pan-KO | 0.9376 | 0.9840 | 0.9945 |

**TABLE V.**
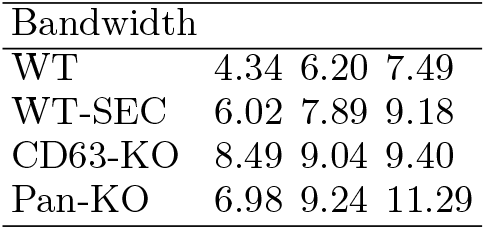
Bandwidth. Bandwidth of the generated PDFs in Fig. 10. Selected with Silverman’s rule of thumb.

**TABLE VI.**
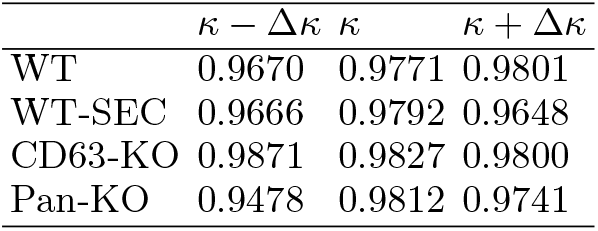
Area under curve. Area under the generated bending modulus probability density functions from the kernel density estimator. The value of 1 indicates a proper probability density function.

| | $\kappa - \Delta\kappa$ | $\kappa$ | $\kappa + \Delta\kappa$ |
| --- | --- | --- | --- |
| WT | 0.9670 | 0.9771 | 0.9801 |
| WT-SEC | 0.9666 | 0.9792 | 0.9648 |
| CD63-KO | 0.9871 | 0.9827 | 0.9800 |
| Pan-KO | 0.9478 | 0.9812 | 0.9741 |

The complete code is available at: https://github.com/thecalculon/Membrane_1D.

#### 3. Are the deformations volume conserving?

Let us try to estimate how well volume conservation is satisfied. The bilipid membrane is permeable to water with a permeability coefficient *p*_w_ which varies from about 10 to 100 × 10^−6^ m/s in bilipid vesicles [76]. The flux of water molecules is given by

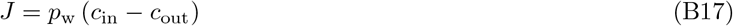

where *c*_in_ and *c*_out_ are the density of water inside and outside the vesicle, respectively. Before the deformation there is no flux, the vesicle is in equilibrium and *c*_in_ = *c*_out_ = *N/V*, where *N* is the number of water molecules inside and *V* is the volume of the vesicle. Now we consider a case where the tip of the AFM moves by a distance *d* in time Δ*t*. The corresponding change in volume is Δ*V* ∝ *d*^3^. The density of water outside remains the same. The density of water inside increases to

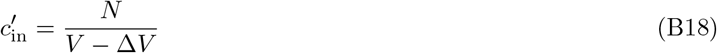

This difference in density drives a flux as given in (B17). The total number of molecules that leave the volume is given by

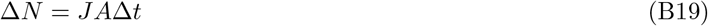

where *A* is the area of the vesicle. We obtain

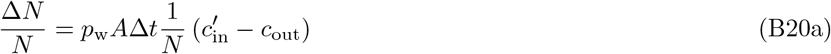

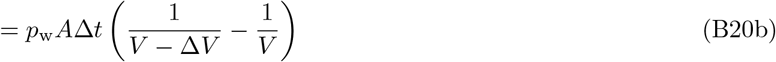

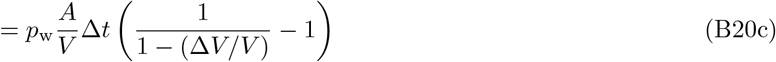

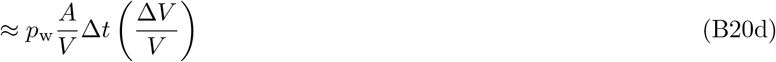

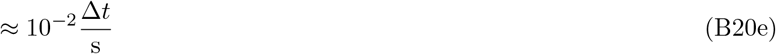

Here we have used *p*_w_ ≈ 1 × 10^−6^ m s^−1^, *R*_v_ ≈ 100 nm, and (Δ*V/V*)sim(*d/R*_v_)^3^ with *d/R*_v_ ≈ 1*/*10. Here we have used 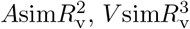 and *d/R*_v_ ≈ (1*/*10) which is certainly true for small deformation. For GUVs *d/R*_v_ lies approximately in the same range as shown in Figure 12.3 in Ref. [13].

## Appendix C Shapiro-Wilk test

## Appendix D PDF of effective elastic modulus

The PDFs were estimated with kernel smoothing function estimate between 0 − 30 mN/m.

## Appendix E PDF of bending modulus

## Appendix F Determination of shape of EVs

## Appendix G Selection rules for force–distance curves

Not all force–distance curves obtained from the indentation measurements were used in the analysis. We describe our selection criteria below.

- The approach and retrace baselines not being constant, not overlapping far away from the vesicle, or having unusually high noise levels. If the approach baseline is not constant the contact point can not be determined correctly, leading to erroneous intervals for determining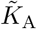.
- During measurements, there is a risk of the AFM drifting or the vesicle moving. Because of this, the curves that show very stiff behavior, comparable to similar measurements made on the substrate, were ignored.
- The curves showing snap-in behavior in the approach were also ignored. While a common trait in air measurements [77], snap-in during liquid measurements could be a sign of tip contamination [6].
- Some vesicles did not show tether formation during retrace. These vesicles were discarded in the extraction of the bending modulus, their approach curves were still used to determine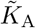.

## Notes

### Competing Interest Statement

The authors have declared no competing interest.

### Summary of Updates

Added additional supporting information parts which are vital to the text.

